# An Artificial Intelligence Framework for Optimal Drug Design

**DOI:** 10.1101/2022.10.29.514379

**Authors:** Grace Ramey, Santiago Vargas, Dinesh De Alwis, Anastassia N. Alexandrova, Joe Distefano, Peter Bloomingdale

**Author notes:** Authors have contributed equally to this work (Grace Ramey, Santiago Vargas).

## Abstract

We introduce the concept of optimal drug design (ODD) as the use of an AI framework to optimize the exposure, safety, and efficacy of drugs. To exemplify the concept of ODD, we developed an artificial intelligence framework that integrates de novo molecular design, quantitative structure activity relationships, and pharmacokinetic-pharmacodynamic modeling. Specifically, our computational architecture has integrated a generative algorithm for small molecule design with a hybrid physiologically-based pharmacokinetic machine learning (PBPK-ML) model, which was applied to generate and optimize drug candidates for enhanced brain exposure. Publicly sourced data on the plasma and brain pharmacokinetics of 77 small molecule drugs in rats was used for model development. We have observed an approximate 30-fold and 120-fold increase on average in predicted brain exposure for AI generated molecules compared to known central nervous system drugs and randomly selected small organic molecules. We believe that with additional data and mechanistic modeling this in silico pipeline could facilitate the discovery of a new wave of optimally designed medicines for the treatment of CNS diseases.

**Graphical Abstract:** Artificial Intelligence Framework for the Optimization of Brain Pharmacokinetics.
A genetic algorithm consisting of cross-breeding, mutating, scoring, and refining was used for de novo generation of a population of new molecular structures. SELFIE representations of molecules were used as input to a variational autoencoder for de novo generation/refinement of individual drug candidates. Molecular descriptors of individual drug candidates are generated and used as input into a trained neural network to generate drug-specific pharmacokinetic (PK) parameters. PK parameters are used as input into a physiologically-based pharmacokinetic (PBPK) model of the brain to predict brain PK of the drug candidate. Brain concentration-time profiles are integrated to obtain an area-under the curve (AUC), a metric of brain exposure, which is used to score and inform the design of new generations of molecules. Iterations of this framework generate novel drug candidates optimized for greater brain exposure. Created with BioRender.

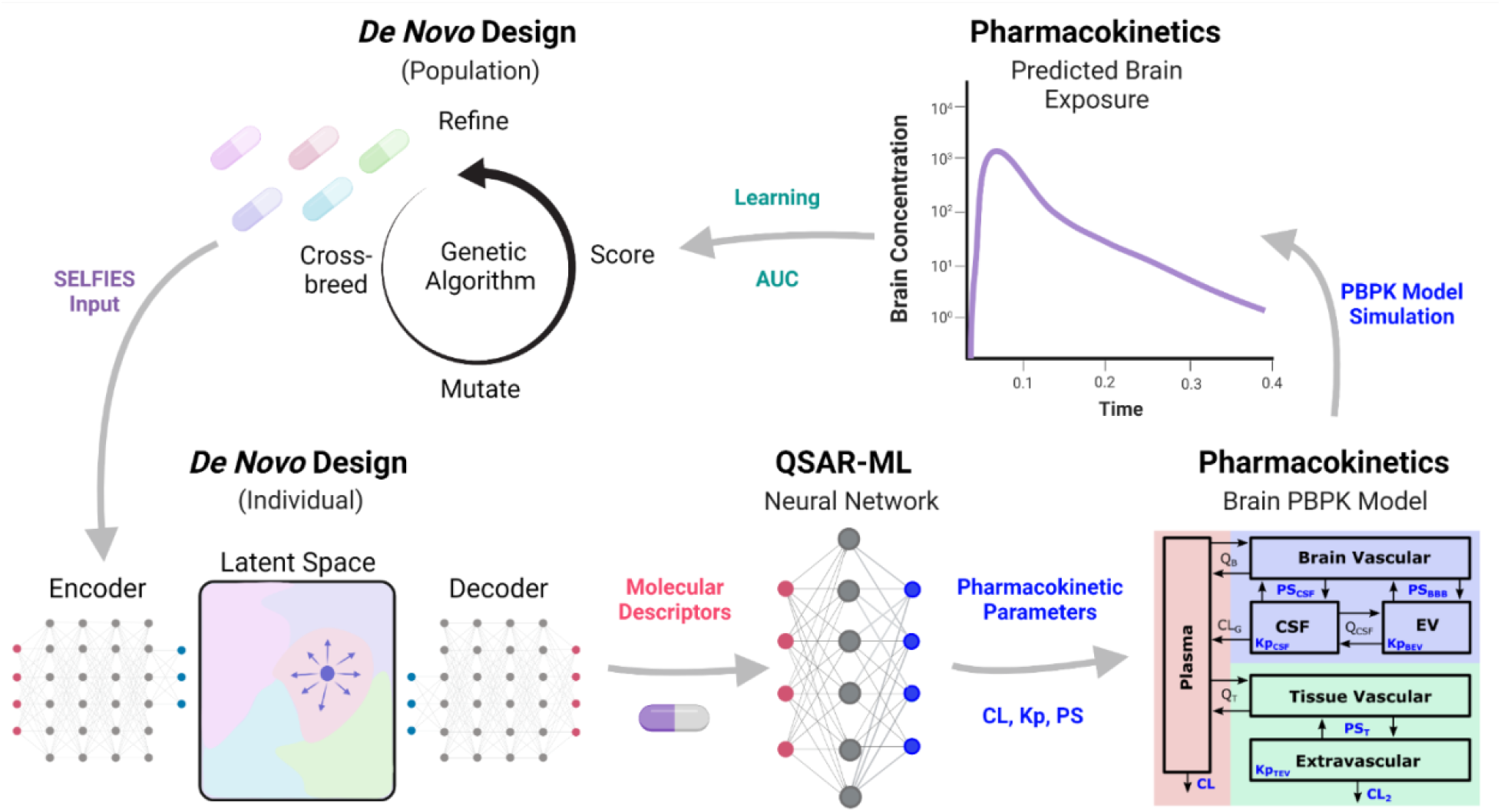

## Introduction

Ever since the introduction of computers into pharmaceutical research and development, computational methods have supported the identification and development of novel drug candidates through molecular design, aided understanding of the quantitative relationship between molecular structure and biological activity, and helped characterize *in vivo* pharmacokinetics/pharmacodynamics (PK/PD)^1,2^. Machine learning (ML), which is anticipated to have a role of increasing importance, has impacted many aspects of drug discovery and development including target and biomarker identification, image analysis workflows, digital clinical endpoints, and drug design and optimization^3^. Pharmaceutical research and development is costly and time consuming and the industry has been under immense pressure to increase productivity^4^. High drug attrition rates continue to burden the industry, especially in neuroscience due to complex pathophysiological mechanisms involved in neurological diseases. Since many pathways are dysregulated in these diseases, lack of efficacy could be due to the focus on single target interventions^5^. Combination therapies or engineering drugs that exhibit a desirable polypharmacological profile could result in efficacious treatment options for unmet neurological diseases. The integration of mechanistic and machine learning models into an artificial intelligence (AI) framework could inform the design and development of new medicines, improving research and development productivity, and overall human health.

Pharmacokinetics (PK) is the time-course of drug absorption, distribution, metabolism, and elimination (ADME) in the body, whereas pharmacodynamics (PD) is the time-course of drug effects on the body. PK-PD modeling is an important component of drug development and aims to provide a quantitative understanding for the relationship between drug exposure and response. Physiologically-based pharmacokinetics (PBPK)^6^ and quantitative systems pharmacology (QSP)^7^ can be viewed as mechanistic extensions of PK/PD, where information about (patho-) physiological systems are used in model structure design and parameter constraints. Although these modeling approaches are commonly applied during preclinical-clinical development, they are seldomly, if ever, implemented in early phases of drug discovery during the design of chemical and biological matter. Quantitative-structure-activity-relationship (QSAR) models are commonly applied during lead candidate optimization and aim to predict an activity or property of a compound using molecular descriptors^8^. Predictions of interest could be metabolic stability, solubility, substrates for drug transporters and metabolizing enzymes, target potency, toxicity and efficacy biomarkers, and many others. QSAR modeling approaches have also been used to correlate molecular descriptors with PK/PD model parameters, referred to as quantitative structure PK/PD relationships (QSPKR/QSPDR), which enables the prediction for the *in vivo* time course of drug exposure and response^9^. Integrating QSAR with PBPK, QSP, and quantitative systems toxicology (QST)^10^ models enables an enhanced quantitative mechanistic understanding of drug exposure, efficacy, and safety, serving as a powerful tool for the design and development of drugs.

De novo molecular design is the automated process of generating novel chemical structures that optimally satisfy a pharmacological profile of interest. The use of AI and ML algorithms in generative models for drug design is very promising, as these algorithms have the potential to revolutionize the drug discovery and development process^11^. The use of AI platforms for drug discovery have already demonstrated the ability to go from initial screening to lead candidate selection and into clinical development in time frames that are much shorter than industry-wide averages^12,13^.

Here we introduce the concept of optimal drug design (ODD) as an AI framework to optimize the exposure, safety, and efficacy of drugs. We exemplify this concept and framework with a case study in which we generate novel molecules optimized for central nervous system (CNS) exposure. Specifically, we developed a fully-integrated *in silico* AI pipeline for ODD by combining algorithms for de novo molecular design, quantitative structure activity relationship (QSAR) modeling, and a brain physiologically-based pharmacokinetic (PBPK) model. We developed and applied our brain PBPK model to estimate PK parameters for drugs. Molecular descriptors were used as features and estimated drug-specific PBPK model parameter values were used as outcomes for the training of machine learning algorithms. This enabled the prediction of drug-specific parameters directly from molecular structure information, which was used as input for the PBPK model to predict plasma and brain PK. Predicted brain exposure was then used to inform the design of new molecular structures. Two types of generative algorithms, a genetic algorithm and variational autoencoder (VAE), were used in combination to generate a population of potential drug molecules and optimize individual molecules. This framework enabled the generation of novel drug candidates and ranked them according to their predicted CNS exposure profile, highlighting this platform’s usefulness for optimal CNS drug design.

## Methods

### Data Collection

Data for plasma and brain pharmacokinetics of small molecular structure compounds in rats and humans was collected from the literature. A comprehensive literature search was conducted in the Embase and PubMed databases. The search strategy included both free-text words and medical subject headings (MeSH), limited to the English language. All records retried from the literature search, using a multi-string search approach, were manually screen based on abstracts and titles. Duplicate records and records that did not match defined eligibility criteria were excluded. Full-text copies for each manuscript were then downloaded and a set of inclusion criteria was applied. Full-text articles were manually screened and available data from the studies were extracted for dataset creation. The dataset contains information related to manuscript details, study details, treatment information (dose, frequency, route of administration, formulation), and outcome details (drug concentrations, sample times, sample size).

The dataset contained brain pharmacokinetic information in rats and humans from 170 publications for 259 compounds. Data was excluded from our analysis based on the following conditions: (a) studies that only measured metabolites, (b) multiple dose data, and (c) compounds with less than three concentration-time data points in plasma and brain. The final datasets we used for PBPK modeling contained 77 and 21 compounds for rats and humans, and compounds that were administered either intravenously or orally as a single dose. Datasets were formatted and processed using R.

### Brain Physiologically-Based Pharmacokinetic Model

We developed a physiologically-based pharmacokinetic model of the brain for small molecule drugs (Fig. 1). The brain PBPK model consisted of six ordinary differential equations and 19 parameters. A minimalistic structure, often referred to as a minimal PBPK (mPBPK) model, was implemented to capture drug exposure in plasma, cerebrospinal fluid (CSF), brain interstitial fluid (ISF), and brain homogenate^14^. The model contains six compartments: plasma, brain vascular, cerebrospinal fluid (CSF), brain extravascular (EV), non-brain tissue vascular, and non-brain tissue EV. There are 12 physiological (Table 1) and 7 (Table 2) drug-specific parameters. Physiological parameter values were obtained from the literature and drug-specific parameter values were estimated for each drug. Physiological parameters include flow rates, compartmental volumes, brain-barrier surface areas, and distributional clearances. In the model, we included fluid flow between plasma and brain vascular (Q_B_), brain extravascular and CSF (Q_CSF_), and plasma and tissue vascular (Q_T_). Compartmental volumes include plasma (V_P_), CSF (V_CSF_), brain vascular (V_Bv_), brain extravascular (V_Bev_), tissue vascular (V_Tv_), and tissue extravascular (V_Tev_). We also incorporated two surface areas to capture drug disposition across the blood-brain-barrier (SA_BBB_) and blood-CSF-barrier (SA_BCSFB_). Glymphatic clearance (CL_G_) was incorporated as a unidirectional flow from CSF to plasma. Drug-specific parameters consist of flow rates across permeable surfaces (PS), partition coefficients (Kp), and elimination clearances (CL). Drugs were able to distribute between brain vascular and CSF at a flow rate of PS_CSF_, brain vascular and brain extravascular space at a flow rate of PS_BBB_, and tissue vascular and tissue extravascular space at a flow rate of PS_T_. To reduce the number of estimated parameters, we estimated the total brain permeable surface (PS_B_) flow rate and scaled this parameter based on the relative brain-barrier surface area to calculate PS_CSF_ and PS_BBB_:

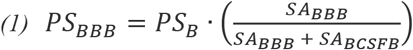

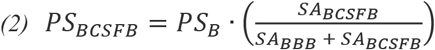

**Fig. 1.**
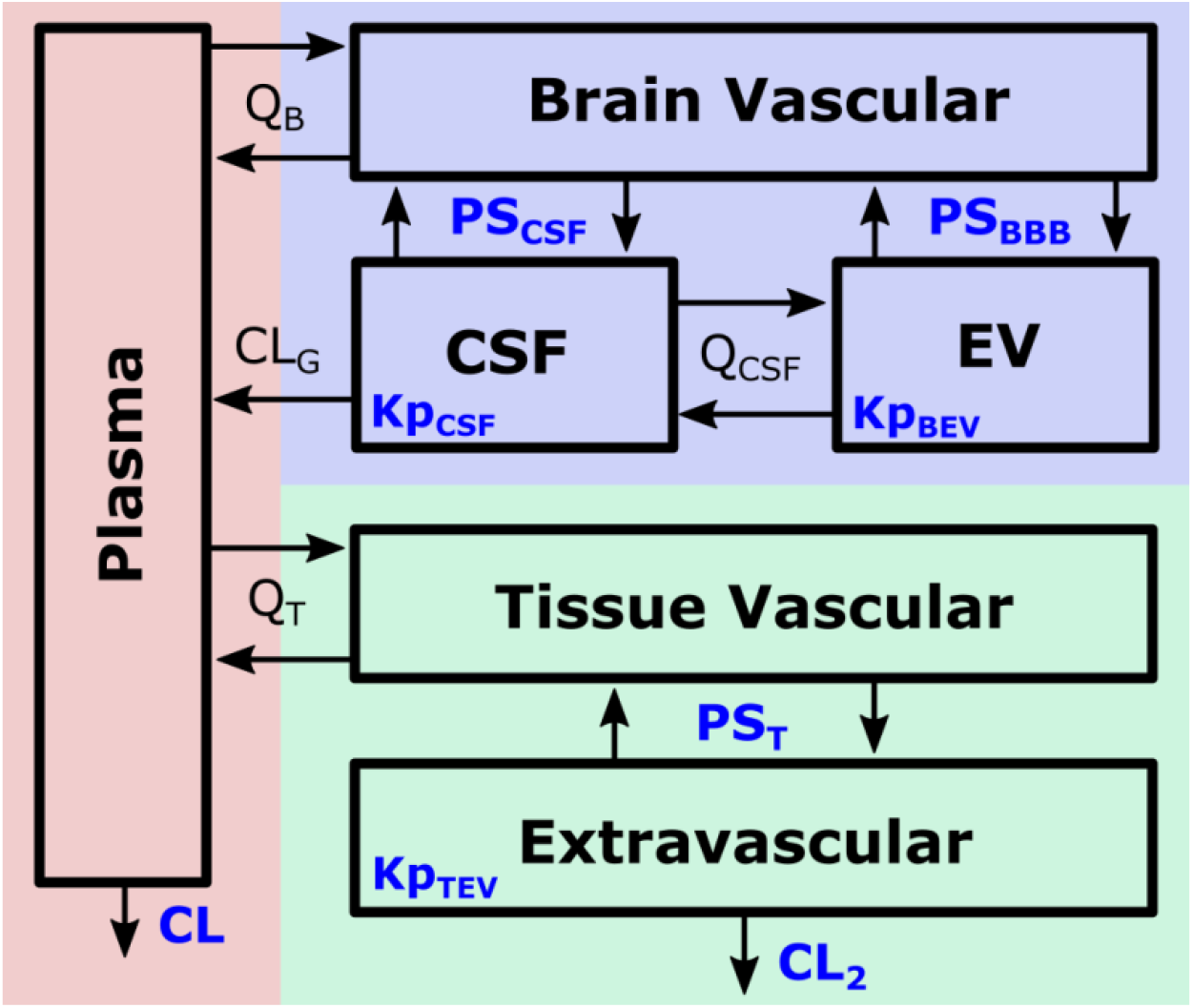
Brain Physiologically-Based Pharmacokinetic (PBPK) Model. A PBPK model of the brain was developed for small molecule drugs, which describes a minimalistic representation of brain anatomical and physiological processes. There are six compartments: plasma, brain vascular, cerebrospinal fluid (CSF), brain extravascular (EV), tissue vascular, and tissue extravascular. The model contains 12 physiological and 8 drug-specific parameters. Four physiological parameters represented here are plasma flow to and from the brain (Q_B_), CSF flow between brain ISF and CSF compartments (Q_CSF_), glymphatic clearance from CSF to plasma (CL_G_), and plasma flow to and from tissues (Q_T_). Additional physiological parameters include compartmental volumes and brain-barrier surface areas. Drug-specific parameters are rate of drug distribution across permeable surfaces (PS), partition coefficients (Kp), and elimination clearances (CL). Drug is able to distribute between brain vascular and CSF at a rate of PS_CSF_, brain vascular and brain extravascular space at a rate of PS_BBB_, and tissue vascular and tissue extravascular space at a rate of PS_T_. Kp_CSF_, Kp_BEV_, and Kp_TEV_, represent partition coefficients between brain CSF/vascular, brain extravascular/vascular, and tissue extravascular/vascular, respectively. There are two elimination clearances processes, from plasma (CL) and tissue (CL_2_). Created with Inkscape.

**Table 1.** Brain PBPK Model: Physiological Parameters in Rats and Humans

| Parameter | Description | Rat | Human |
| --- | --- | --- | --- |
| $V_P$ | Plasma volume | 9.06 mL | 3.13 L |
| $V_B$ | Total brain volume | 2.28 mL | 1.45 L |
| $V_{Bv}$ | Brain vasculature volume | 0.0502 mL | 0.0319 L |
| $V_{Bev}$ | Brain extravasculature volume | 2.11 mL | 1.27 L |
| $V_{CSF}$ | Cerebrospinal fluid volume | 0.120 mL | 0.150 L |
| $V_T$ | Total non-brain tissues volume | 269 mL | 69.6 L |
| $V_{Tv}$ | Non-brain tissues vasculature volume | 7.89 mL | 1.68 L |
| $V_{Tev}$ | Non-brain tissues extravasculature volume | 261 mL | 64.7 L |
| $Q_B$ | Plasma flow rate to brain | 65.3 mL/h | 21.5 L/h |
| $Q_T$ | Plasma flow rate to non-brain tissues | 2880 mL/h | 160. L/h |
| $Q_{CSF}$ | CSF flow rate between ventricles and brain extravasculature | 0.132 mL/h | 0.0100 L/h |
| $CL_G$ | Glymphatic clearance | 0.120 mL/h | 0.0230 L/h |
| $SA_{BBB}$ | Surface area of the blood-brain-barrier | 0.0155 m<sup>2</sup> | 20.0 m<sup>2</sup> |
| $SA_{BCSFB}$ | Surface area of the blood-cerebrospinal-fluid-barrier | 0.00250 m<sup>2</sup> | 7.5 m<sup>2</sup> |

**Table 2.**
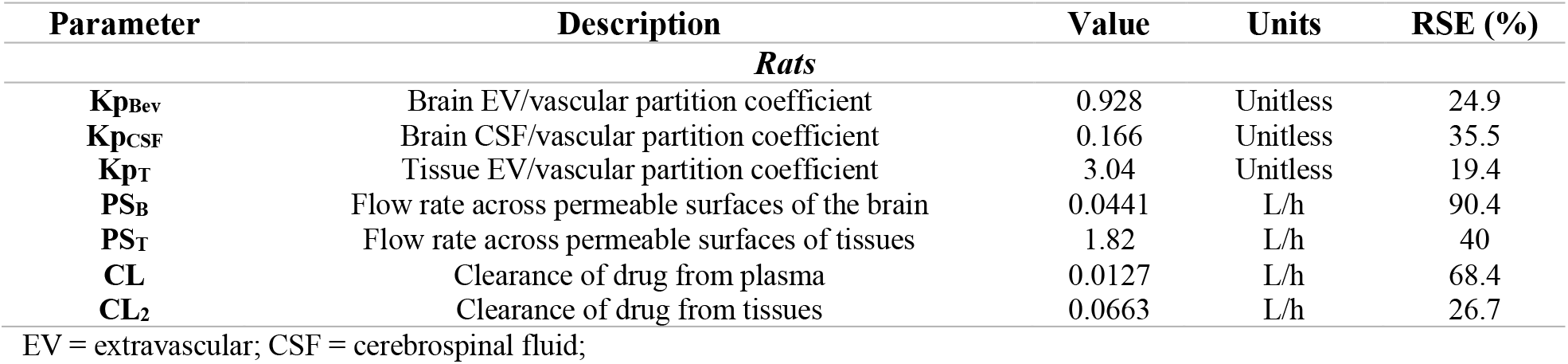
Brain PBPK Model: Drug-Specific Parameters (Population Estimates) in Rats

We included partition coefficients for capturing concentration differences between brain CSF/vascular (Kp_CSF_), brain extravascular/vascular (Kp_Bev_), and tissue extravascular/vascular (Kp_Tev_). Lastly, there were elimination clearances from both plasma (CL) and non-brain tissues (CL_2_). Brain PBPK model equations are:

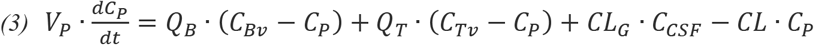

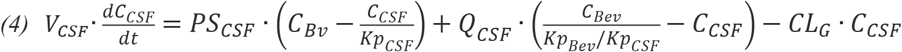

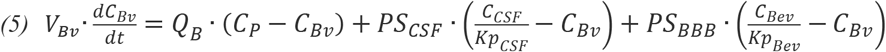

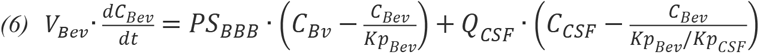

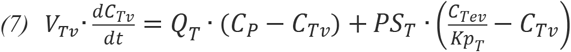

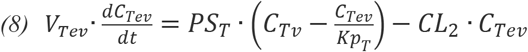

Where, C_P_, C_CSF_, C_Bv_, C_Bev_, C_Tv_, and C_Tev_, are the concentrations of drug in plasma, CSF, brain vasculature, brain extravasculature, tissue vasculature, and tissue extravasculature, respectively.

Nonlinear mixed-effect modeling was conducted in Monolix (version 2019R2) using the brain PBPK model and PK data described above. A population-based approach was implemented to enable a simultaneous estimation for 77 drugs in rats, where each drug essentially acts as a separate individual. The estimation of population parameters was performed using stochastic approximation expectation-maximization algorithm (SAEM). Estimated drug-specific parameters for the entire population of drugs are described in Table 2 and individual parameter values for each drug are described in Supplementary Table 1. RSE in population parameter estimates, −2 log likelihood (−2LL), Akaike information criteria (AIC), and Bayesian information criteria (BIC), were used for model selection criteria. Observed versus predicted plots for drug concentrations were generated for plasma, CSF, brain ISF, and brain homogenate. Brain homogenate was determined based on concentration of drug in brain extravascular and CSF ventricles, which assumes that brain vasculature was perfused prior to collection and homogenization. The brain EV compartment represents concentration of drug in both brain ISF and intracellular fluid, which assumes that the concentration of drug in brain ISF is equivalent to intracellular concentrations.

### QSAR-Based Machine Learning Models

Machine learning models were developed to predict PBPK model parameters using quantitative molecular structure information. Simplified molecular-input line-entry system (SMILES) strings were used to generate molecular descriptors using Mordred and RDKit^15,16^. Generated molecular descriptors for each drug were used as input to two different machine learning algorithms and estimated drug-specific PBPK model parameter values were used as output. Support vector regression (SVR) models, a class of support vector machines, were used as they allow for fitting of more complex data in higher dimensional hyperspace, which is a good option for linking both continuous and discrete molecular descriptors to parameter values. Multilayer perceptron (MLP) models, a class of artificial neural networks, were also used, which consists of one hidden layer and heavy regularization to avoid overfitting. Five-fold testing was used to select a set of hyperparameters. Transfer learning was evaluated to see if model performance was improved by imputing predicted values for one parameter into the model for another, but no appreciable improvements were made. An 80/20 split of the data was used for the training/test sets.

### De Novo Molecular Design Algorithms

SELF-referencIng Embedded String (SELFIES), a recent advancement beyond SMILES, representation of molecules has been shown to generate molecules with greater molecular validity^17^. We have implemented the combination of a genetic algorithm and variational autoencoder (VAE) that use SELFIES molecular representations to generate de novo molecules. A genetic algorithm was created from scratch to generate a population of molecules through four steps: mutate, cross-breed, score, and refine. A mutation rate of 0.1 and random crossover events were used to create offspring. Molecules were scored based on their predicted brain exposure (AUC) and filtered based on favourable Lipinski criteria. For this work, two or more of the five Lipinski rules must be satisfied for a compound to be considered pharmaceutically viable. Additionally, we have implemented VAEs to map molecular structures generated with the genetic algorithm to a latent space. This approach allows for the use of gradient based optimization and has been explored as a route to optimize molecular function by others^18^. The VAE was trained using filtered data from ChEMBL and the QM9 dataset^19,20^. Gradient optimization within this latent space better interpolates between molecules and enables the generation of molecular structure variations relative to a point of origin. The advantage of these algorithms is that they allow for the optimization of individual molecules rather than populations of molecules. In other words, the genetic algorithm enabled a broad generation of diverse chemical matter, whereas the VAE enabled the fine-tuning of molecular structures. Model architecture is depicted in Supplementary Table 3.

### Proof-of-Concept for Drug Candidates Optimized for Enhanced Brain Exposure

Using our platform, we generated 300 new molecules and compared their predicted brain exposure (AUC) to ~300 molecules in the QM9 dataset and ~300 CNS drugs and drug candidates obtained from the literature. We have considered molecules in the QM9 dataset as a control as we expected the newly generate molecules to display improved brain exposure compared to randomly select organic molecules. The PK of each molecule was simulated using the same dose and brain AUCs were determined over 1 day (AUC_0-24h_).

## Results

### Characterization of CNS PK using a hybrid brain PBPK machine learning model

We developed a PBPK model of the brain for small molecule drugs (Fig. 1) and used it to characterize the plasma and CNS PK of 77 drugs in rats. The model consists of six equations and 19 parameters, described in detail in methods. There were 12 physiological parameters (Table 1) in the model that remained fixed during the estimation of 7 drug-specific parameters (Table 2). A population-based modeling approach was used to enable the simultaneous estimation of pharmacokinetic parameters across all drugs. Therefore, in the estimation of drug-specific parameters, a population mean estimate was obtained for all drugs as well as individual estimates for each drug. Five of the seven estimated population mean parameters had percent relative standard errors (RSE) values < 50%, while PS_B_ and PS_T_ were estimated with percent RSE values of 90.4% and 68.4%, respectively. Percent RSE for random effects were all < 35% (Supplementary Table 2). The low percent RSE values for the parameters as well as the observed versus predicted plots for drug concentrations in rat plasma, brain homogenate, brain ISF, and CSF (Fig. 2) indicated that the model characterizes PK in rats well.

**Fig. 2.**
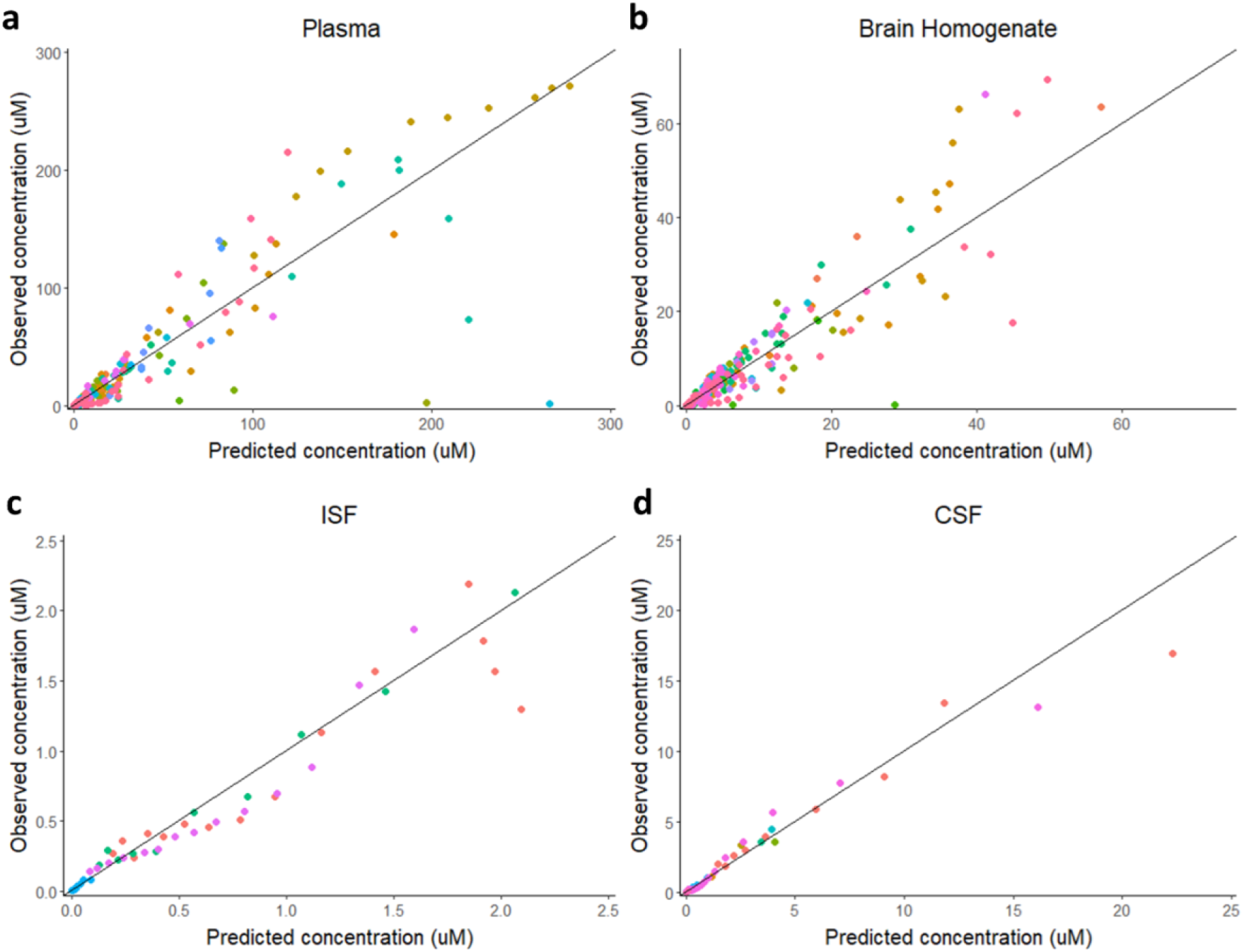
Observed vs Predicted Drug Concentrations in Rats. Observed experimental data for the concentration of 77 drugs along with PBPK model predictions are shown for (a) plasma, (b) brain homogenate, (c) brain ISF, and (d) CSF. The black solid line represents the line of identity. Colors represent different drugs.

We used molecular descriptors and PBPK model estimated drug-specific parameter values to train QSAR-based machine learning models. Molecular descriptors were generated from the molecular structure for each compound in the dataset. Molecular descriptors and their relative importance for the prediction of each drug-specific parameter are described in Supplementary Materials 1. We used single regression machine learning models to make drug-specific parameter predictions, the results of which are listed in Table 3. Support vector regression (SVR) models were the best for predicting Kp_Bev_, Kp_CSF_, PS_T_, and CL_2_, yielding R^2^ values of 0.42, 0.20, 0.20, and 0.29. The number of molecular descriptors selected based on feature importance and used to inform predictions of these parameters were 40, 50, 100, and 60, respectively. Multilayer perceptrons (MLPs) yielded the best predictions for Kp_T_, PS_B_, and CL, with R^2^ values of 0.19, 0.33, and 0.25. The number of features selected for the predictions of these parameters were 10, 20, and 50, respectively.

**Table 3.** Single Regression Machine Learning Results for Drug-Specific Parameters

| Parameter | Validation Performance ($R^2$) | Features (#) | Algorithm |
| --- | --- | --- | --- |
| $K_{pBev}$ | 0.42 | 40 | Linear SVR |
| $K_{pCSF}$ | 0.20 | 50 | Linear SVR |
| $K_{pT}$ | 0.19 | 10 | MLP |
| $PS_B$ | 0.33 | 20 | MLP |
| $PS_T$ | 0.20 | 100 | Linear SVR |
| $CL$ | 0.25 | 50 | MLP |
| $CL_2$ | 0.29 | 60 | Linear SVR |

SVR = support vector regression; MLP = multilayer perceptron;

Subsequent to model development and training, QSAR-based machine learning algorithms were integrated with PBPK model equations into a single pipeline to enable the generation of plasma and brain concentration-time profiles directly from molecular structure.

### Generation of novel drug candidates optimized for brain exposure

We combined a genetic algorithm and VAE for the de novo generation of molecules. De novo design algorithms were set up as input into the hybrid brain PBPK-ML model, which enabled the prediction of brain concentration over time for each newly generated molecule. The genetic algorithm was used to simulate a population of 1000 molecules. Upon each new generation of molecules, the original population of 1000 molecules were mutated, cross-bred, scored, and refined to arrive at molecules that were predicted to have improved brain exposure. An average and median improvement in brain area under the concentration-time curve (AUC) from the first to fifth generation of 2-fold and 6-fold for the top 100 molecules was achieved using the genetic algorithm (Fig. 3a). Individual molecules were subsequently modified using a VAE and a gradient-based optimization algorithm to further improve predicted brain exposure. An average and median improvement in brain AUC from the seed to final molecules of 46-fold and 9-fold was achieved using the VAE architecture (Fig. 3b). To exemplify the change in brain exposure as molecules traverse the VAE latent space, we depict three different trajectories of molecule (Fig. 3c). We then aimed to provide an *in silico* proof of concept for the generation of novel drug candidates that are optimized for improved brain exposure. We compared the predicted brain exposure for molecules in the QM9 dataset (set of random small organic molecules), CNS drugs and drug candidates gathered from the literature, and AI designed molecules using the platform we have developed. There was approximately a 30-fold and 120-fold increase on average in predicted brain exposure for the AI generated molecules optimized on brain exposure compared to existing CNS drugs and QM9 groups, respectively (Fig. 4).

**Fig. 3.**
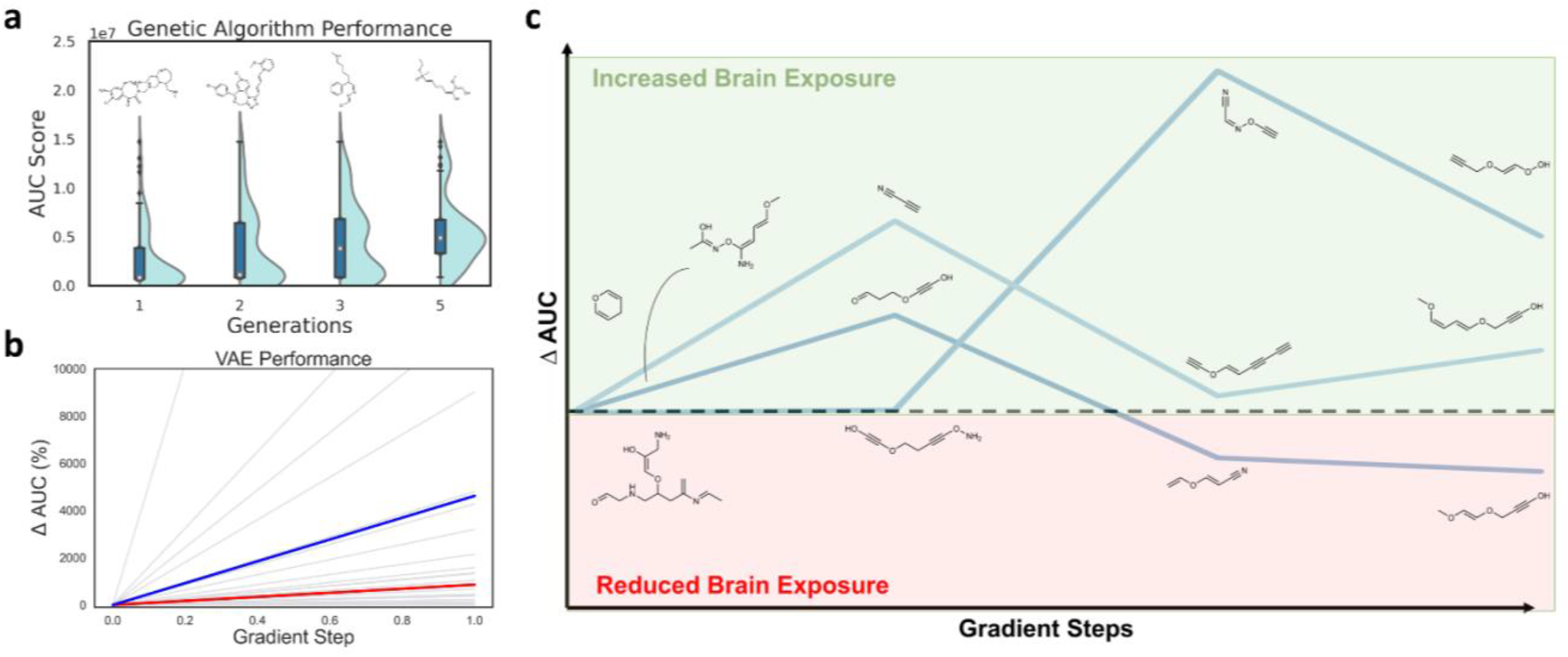
Generative Algorithm and VAE Performance. (a) The change in the mean and distribution in predicted brain homogenate exposure (AUC) for the top 100 performing molecules in a population of 1000 molecules per generation across several generations of the genetic algorithm. The top performing molecule in each generation is displayed. (b) Change in an individual molecule predicted brain homogenate exposure (Δ AUC percent) over the course of a short VAE gradient ascent run. Every step along the trajectory of 10 gradient update steps is not represented, only the change in AUC at the start and end of the run is depicted. Individual, mean, and median changes in AUC are shown as gray, blue, and red lines. (c) An example for the change in predicted brain exposure of three trajectories of a molecule over three gradient update steps.

**Fig. 4.**
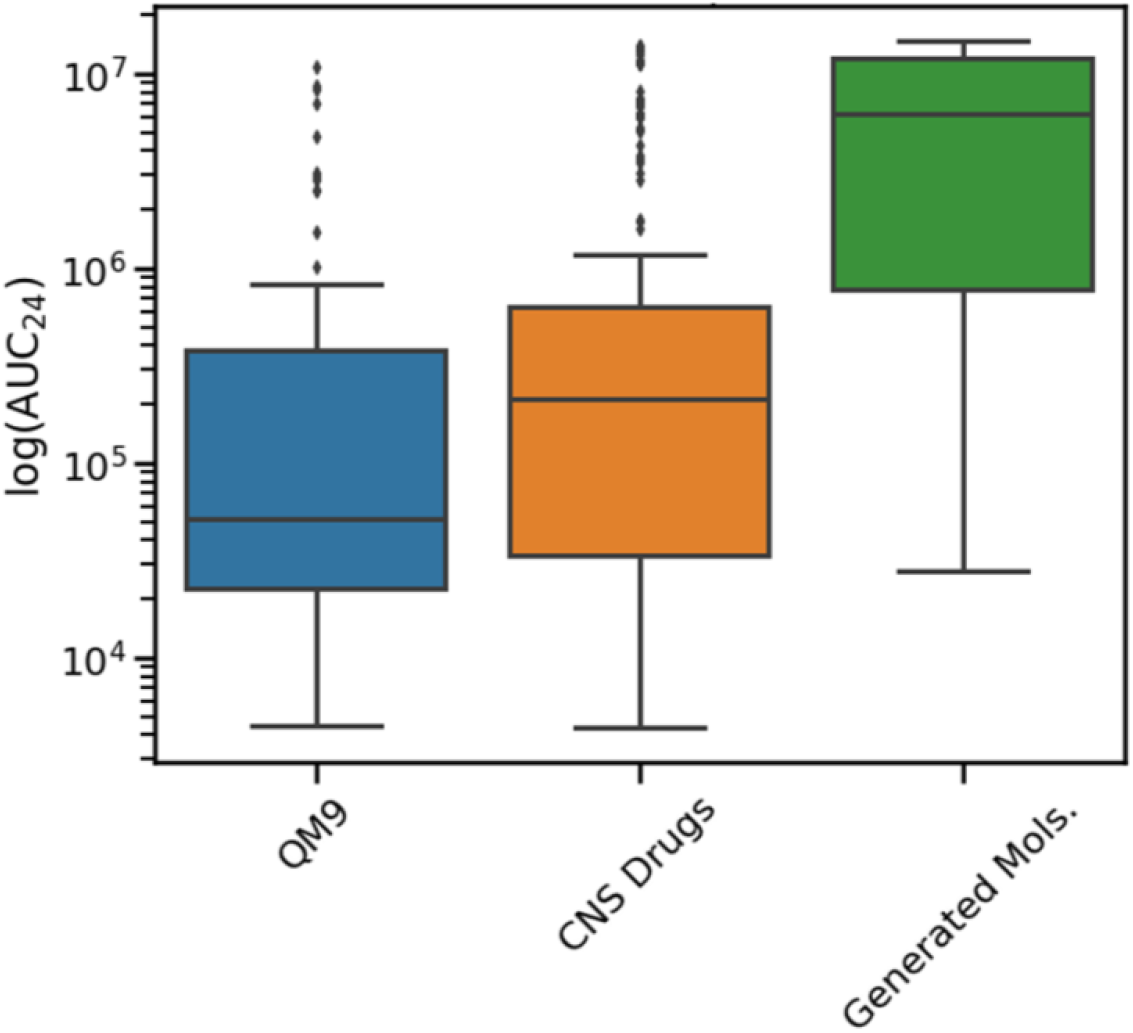
Generation of Novel Molecules with Optimized Brain Pharmacokinetics. Comparison of the predicted brain exposure (AUC) for drugs in the QM9 dataset, CNS drugs dataset, and de novo generated molecules. There are approximately 300 drugs evaluated in each group. QM9 (blue) is a standard dataset of small organic molecules, utilized here as a random set of molecules. CNS drugs (orange) dataset is from our literature search for compounds that have been investigated for their CNS exposure in animal and humans. Generated mols (green) are molecules that were generated using the modeling framework we have developed.

## Discussion

To our knowledge, we have developed the first fully integrated platform for optimal drug design. There are many examples for the de novo generation of leads in pharmacology using reinforcement learning, generative adversarial networks (GANs), and variational autoencoders^21^. However, none of these prior approaches have incorporated pharmacometrics modeling. We have integrated generative and evolutionary algorithms, QSPKR-based machine learning, and PBPK modeling, which allowed us to generate novel molecules optimized for improved brain exposure. We utilized a population-based approach, nonlinear mixed-effect modeling, to simultaneously model the plasma and brain PK of 77 drugs in rats. Parameter estimates and molecular descriptors for each drug were used in the training of QSPKR models. A novel de novo molecular design method consisting of a genetic algorithm and VAE was used to identify a group of newly generated molecules optimized for greater brain exposure.

QSPKR-QSPDR models have not yet been commonly adopted in practice. QSAR models employed for multi-parameter optimization in drug discovery have focused primarily on predictions of in vitro ADME properties, target binding, and toxicity and efficacy biomarkers. QSAR models have been applied to predict ADME properties (e.g., transporter binding, metabolic stability, efflux ratios, tissue-to-plasma concentration ratio, mean residence time, protein binding, etc.) that can be associated to *in vivo* PK, while QSPKR models directly enable *in vivo* PK predictions. Therefore, lead candidates could be ranked based on *in vivo* PK, rather than a combination of various ADME properties.

Depending on the level of mechanistic detail included in the model, there may be many processes convoluted into a single drug-specific parameter. For example, in our brain PBPK model there are multiple processes that govern the rate of drug uptake into the brain (PS_B_), such as passive diffusion and transporter uptake and efflux. Future expansions of this model could consider transporter efflux (e.g., p-glycoprotein and BCRP), which could be implemented mechanistically into the PBPK model or empirically by using *in vitro* measurements of transporter binding or efflux ratios as a feature in the QSPKR model for the PS_B_ parameter.

A major limitation of our analysis is the sparsity and quality of data. The number of compounds (77) used in this analysis is insufficient for robust development of a hybrid PBPK-ML model. For each compound, there was also a limited number of time points for CNS concentrations. Since our dataset was created through the pooling of publicly available data in the literature, the quality and consistency of data between different labs is of concern. Inter-lab differences in methodologies for the determination of plasma and brain drug concentrations could introduce a greater level of variability. Although, the majority of population parameter estimates had percent RSE values <50%, which indicated a reasonably acceptable level of confidence, two parameters (PS_B_ and PS_T_) had percent RSE values outside of a desirable range (>50%). Test set R^2^ values for the relationships between drug-specific parameters and molecular information ranged from 0.19 to 0.42, with an average of 0.27, which indicates low predictive accuracy. To note, the primary goal of our research was to fully-integrate multiple model platforms into a single AI framework to highlight the applicability for ODD, rather than provide a strong validation of model performance. We believe that the models would have improved performance with the inclusion of more data as additional datasets arise in the future. A consideration to increase throughput and decrease variability of preclinical PK studies and to potentially improve preclinical-clinical translatability is the use of microfluidic organ-on-a-chip technologies ^22^.

Another limitation is the limited availability of brain PK in humans. We have collected data from clinical studies for 21 drugs in humans, which overlap with drugs in our rat dataset, but most of these studies have only measured plasma PK. This poses a difficult problem, because the primary objective is to optimize the design of drugs for humans, not preclinical animal species. To overcome the lack of clinical data, one consideration could be to use allometric relations between rats and humans, which is a common methodology for human dose predictions. Alternatively, in the absence of data, transfer learning methods between rat and human QSPKR models could be explored. We have attempted to fit the PBPK model to the human data, but parameter estimates had a significantly high level of uncertainty. Additional data is required to increase the reliability of predicting drug exposure in human brain.

Despite limitations in data availability, we were able to develop and integrate a set of models/algorithms that generated novel molecules predicted to exhibit enhanced brain exposure in rats. Brain exposure of newly generated molecules were compared against a random population of small organic compounds and a druglike compounds from the ChEMBL dataset. Generated molecules exhibited orders of magnitude increases in predicted brain exposure, relative to existing CNS drugs and druglike molecules. The combination of a genetic algorithm and VAE provided a powerful means for high-throughput molecular generation as well as a focused refinement. Trends from these algorithms are also in agreement with previous findings that more lipophilic molecules with rigid structures and lower polar surface area tended to bypass the brain-blood barrier with greater ease^23^. We observed that total polar surface area and logP increased for both the genetic and variational autoencoder schemes relative to the starting set of molecules.

In this research we focused solely on optimizing for improved brain exposure; however, multi-objective optimization could also be performed when there are multiple endpoints of interest. Endpoints could consist of *in silico* parameters or measurements from *in vitro/vivo* experiments. Multiple mechanistic models, such as PBPK, QSP, and QST, could be all be leveraged for the simultaneous optimization of drug exposure, efficacy, and safety. These models should be viewed as modular, where they can be swapped in/out depending upon the drug, system, or property of interest. We believe that this work provides a proof-of-concept for the potential of AI frameworks to optimize the design of drugs and a vision for how end-to-end in silico pipelines could operate in the future. Ultimately, streamlining the drug development process and creating novel first/best-in-class medicines.

## Study Highlights

### What is the current knowledge on the topic?

The potential for artificial intelligence and machine learning to revolutionize drug discovery and development is often touted, but original research on this topic is seldom as the field continues to explore novel applications.

### What question did this study address?

The primary question was to understand the technical feasibility for integrating multiple modeling components as well as to assess if the platform is generating molecules that are predicted to have greater brain exposures. Proof of concept of an artificial intelligence framework for drug optimization.

### What does this study add to our knowledge?

This work provides an in silico proof-of-concept for the application of an artificial intelligence framework to optimize the structure of small molecule drugs. De novo molecular design, QSAR, and brain PBPK models were integrated to discover novel molecules optimized for enhanced brain exposure. This work also highlights data and model architecture prerequisites for the adoption of artificial intelligence frameworks.

### How might this change drug discovery, development, and/or therapeutics?

The methodology presented could act as an alternative to or augment the current drug discovery and development process. The integration of various machine learning and mechanistic models into a single framework enables the simultaneous optimization of drug exposure, safety, and efficacy. We envision the adoption of fully integrated artificial intelligence frameworks embedded in early drug discovery programs that are continuously trained on emerging high-quality preclinical and clinical data.

## Acknowledgements

We would like to acknowledge Ka Lai Yee and Robert Sheridan for their mentorship and guidance

## Author Contributions

GR, SV, and PB wrote manuscript, designed research, performed research, and analyzed data. DDA, AA, and JD contributed to the research design and manuscript revisions.

## Supplementary Information

Supplementary Materials 1: Molecular descriptors and their relative importance for the prediction of each drug-specific parameter are described

Supplementary Materials 2: Excel sheet containing Monolix formatted dataset of drug concentrations

Supplementary Materials 3: Monolix brain PBPK model text file

Supplementary Materials 4: Excel sheet containing individual drug-specific parameters, random effects, and residual errors, and VAE Model Architecture

## Data and Code Availability

PBPK model code is available in Supplementary Material 3 and code related to computational chemistry algorithms can be found at https://github.com/alexandrova-lab-ucla. The dataset used for PBPK modeling of plasma and brain drug concentrations in rats is available in Supplementary Material 2.

