## Supplementary Materials 1 for "An Artificial Intelligence Framework for Optimal Drug Design"

**Example SMILES String of Aspirin:** CC(=O)OC1=CC=CC=C1C(=O)O

**Example Descriptors:**

**Structural:**

**BIC[x]:** [x]-ordered bonding information content
**C[x]SP[y]:** SP[y] carbon bound to [x] other carbons
**Morg[x]:** 3D-MoRSE (distance = [x])
**n6Ring:** # of 6-member rings
**nAcid:** # of acidic groups
**nBase:** # of basic groups
**nBondsD:** # of double bonds
**NdO:** # of dO
**NdssC:** # of dssC
**Nspiro:** # of spiro atoms
**NtsC:** # of tsC
**SaaCH:** # of aaCH
**SddssS:** # of ddssS
**SsssB:** # of sssB
**SIC[x]:** [x]-ordered structural information content
**ZMIC[x]:** [x]-ordered Z-modified information content

**Electronic:**

**AATS[x][are/c/dv/i/m/p/pe/se/Z]:** averaged and centered moreau-broto autocorrelation of lag [x] weighted by allred-rocow electronegativity/gasteiger charge/valence elections/ionization potrential/mass/polarizability/pauling electronegativity/sanderson electronegativity/atomic number]

**ATS[x][are/c/dv/i/m/p/pe/se/Z]:** moreau-broto autocorrelation of lag [x] weighted by [allred-rocow electronegativity/gasteiger charge/valence elections/ionization potrential/mass/polarizability/pauling electronegativity/sanderson electronegativity/atomic number]

**ATSC[x][are/c/dv/i/m/p/pe/se/Z]:** centered moreau-broto autocorrelation of lag [x] weighted by [allred-rocow electronegativity/gasteiger charge/valence elections/ionization potrential/mass/polarizability/pauling electronegativity/sanderson electronegativity/atomic number]

**GATS[x][are/c/dv/i/m/p/pe/se/Z]:** geary coefficient of lag [x] weighted by [allred-rocow electronegativity/gasteiger charge/valence elections/ionization potrential/mass/polarizability/pauling electronegativity/sanderson electronegativity/atomic number]

**MATS[x][are/c/dv/i/m/p/pe/se/Z]:** moran coefficient of lag [x] weighted by [allred-rocow electronegativity/gasteiger charge/valence elections/ionization potrential/mass/polarizability/pauling electronegativity/sanderson electronegativity/atomic number]

**PEOE_VSA[x]:** MOE Charge VSA Descriptor [x]
**SlogP_VSA[x]:** MOE logP VSA Descriptor [x]
**SMR_VSA[x]:** MOE MR VSA Descriptor [x]
**VSA_Estate[x]:** [x]th EState VSA
**Xc-[x][d/dv]:** [x]-ordered Chi cluster weighted by sigma/valence electrons
**Xch-[x][d/dv]:** [x]-ordered Chi chain weighted by sigma/valence electrons
**Xpc-[x][d/dv]:** [x]-ordered Chi path-cluster weighted by sigma/valence electrons
**GGI[x]:** [x]-ordered raw topological charge

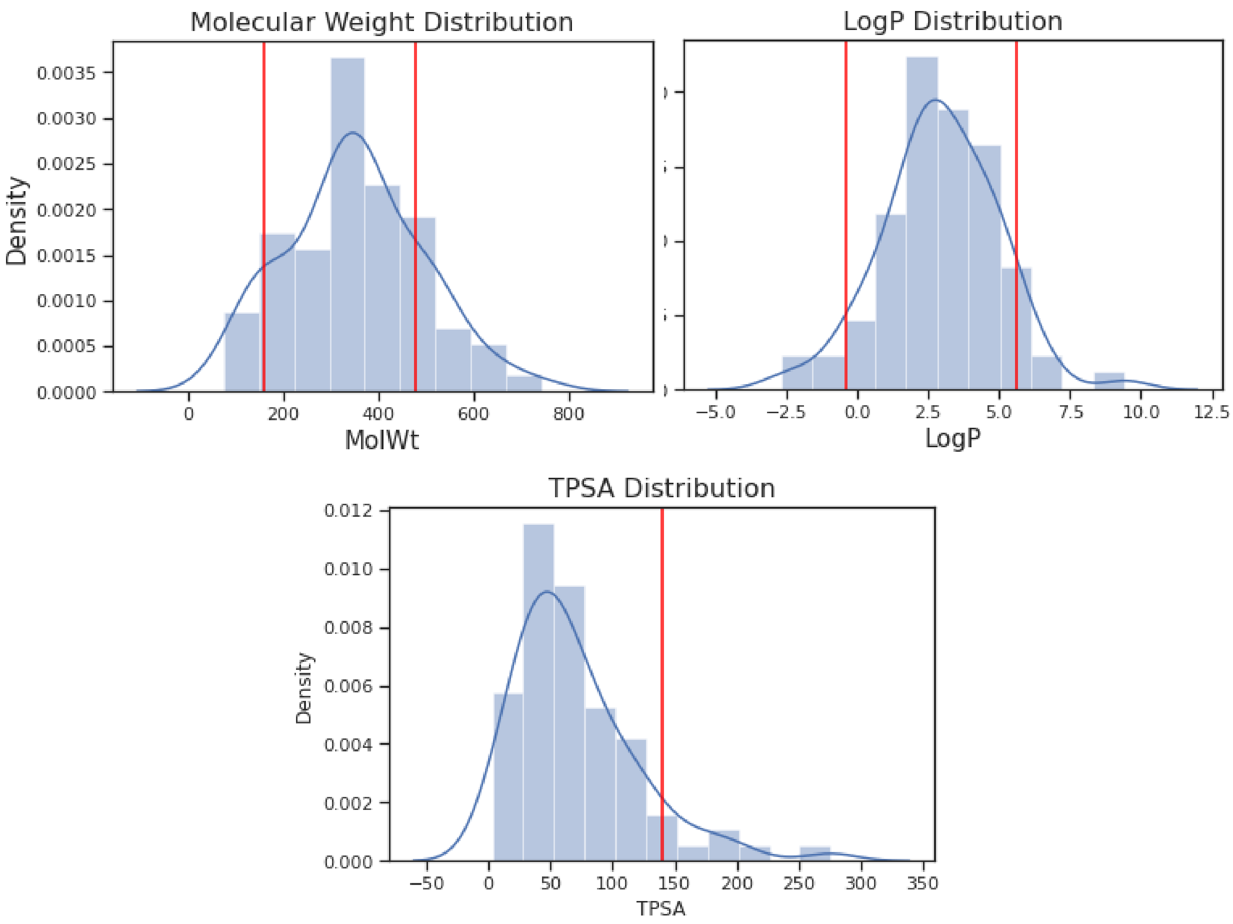


The distribution of our filtered PK dataset for various important benchmark parameters including A) molecular weight (MolWt), B) log water-octanol partition (logP), and C) topological polar surface area (TPSA). The red lines denote filter values for Ghose[] and Veber[] druglike molecular filters. Most of our dataset fell within often used boundaries for all three of these values, and most of our training molecules were small molecules (160 < MolWt < 480) with limited fatty character (-0.4 < LogP < 5.6).

### $K_{pCSF}$

**Structural:** SddssS, nBondsD, NdO, nAcid, ZMIC4
**Electronic:** GGI1, ATS4m, ATS4dv, ATS5m, ATS5dv, ATS6m, Xc-4d, Xc-3d, Xpc-4d

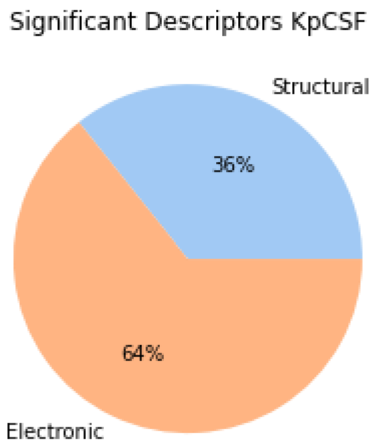


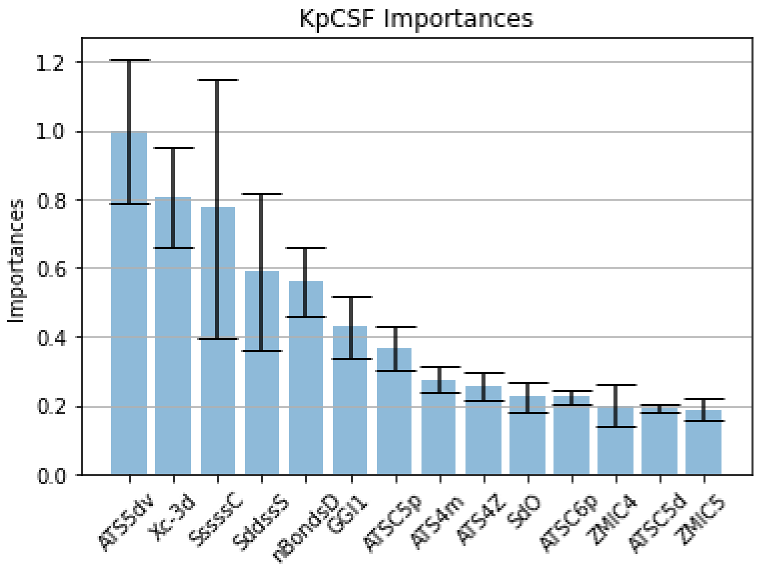


| Variable | P-score | F-score |
|---|---|---|
| Xc-3d | 0.002 | 11.103 |
| nBondsD | 0.004 | 9.543 |
| GGI1 | 0.006 | 8.914 |
| Xc-4d | 0.006 | 8.8 |
| ATS4dv | 0.008 | 8.241 |
| ZMIC5 | 0.008 | 8.021 |
| Xpc-4d | 0.008 | 7.996 |
| ATS4m | 0.01 | 7.603 |
| ATS5dv | 0.01 | 7.522 |
| NdO | 0.011 | 7.456 |
| ZMIC4 | 0.011 | 7.323 |
| ATS3m | 0.011 | 7.426 |
| nAcid | 0.013 | 7.034 |
| ATS5m | 0.014 | 6.912 |
| ATS6m | 0.015 | 6.747 |
| ATS4Z | 0.015 | 6.748 |

#### $Kp_{Bev}$ ($K_{pISF}$)

**Structural:** nBondsD, nAcid, NdssC
**Electronic:** ATSC0se, ATSC0pe, ATSC2Z, Xch-5dv, PEOE_VSA5, ATSC5c

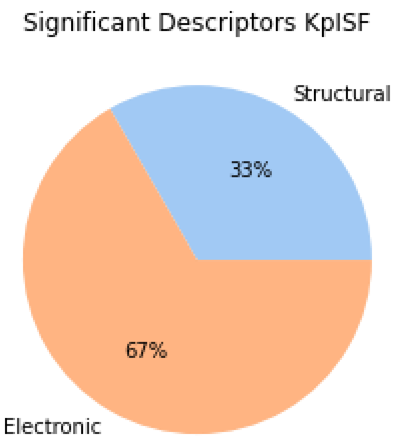


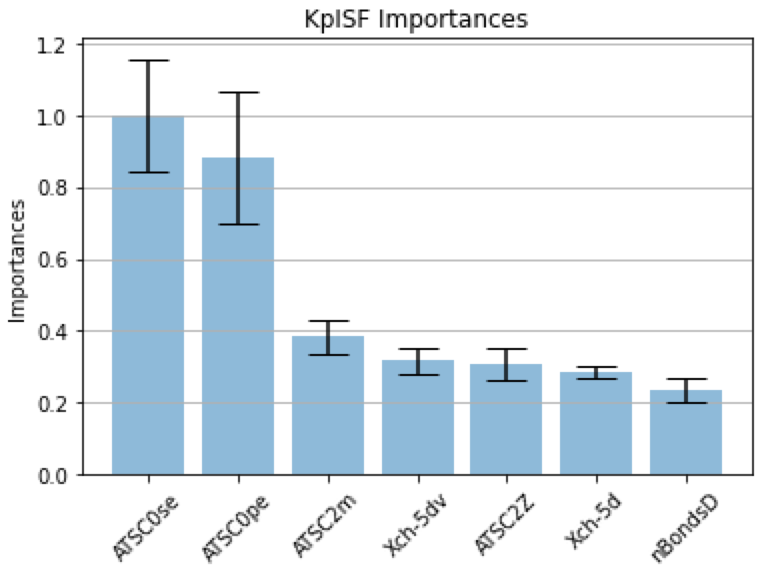


| Variable | P-score | F-score |
|---|---|---|
| ATSC5c | 0.021 | 5.774 |
| PEOE_VSA5 | 0.017 | 6.151 |

# $K_{pT}$

**Structural:** SaaCH, n6Ring
**Electronic:** AATSC0are, AATSC0pe, AATSC0se, SMR_VSA1, EState_VSA10, SlogP_VSA6, VSA_EState6

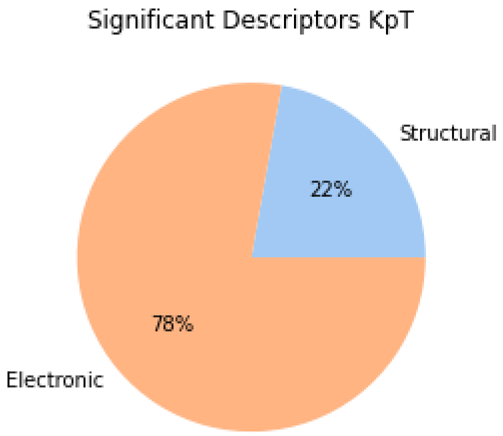


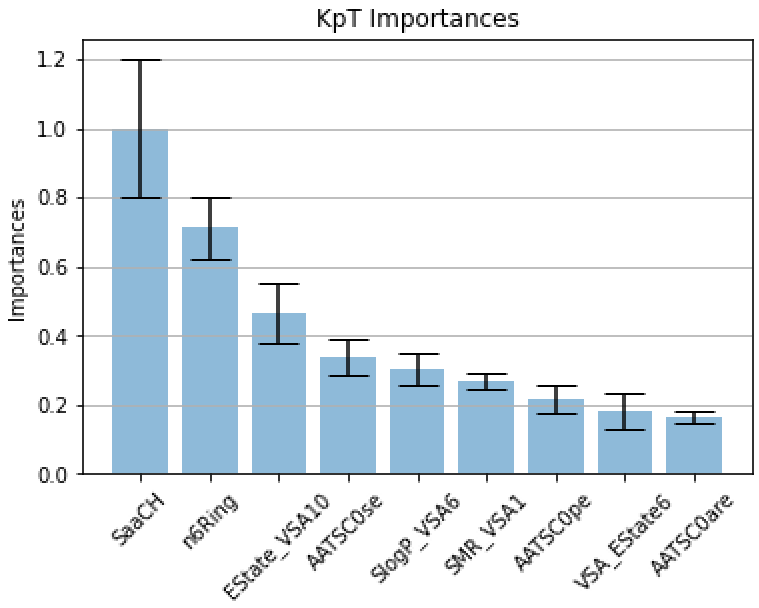


| Variable | P-score | F-score |
|---|---|---|
| n6Ring | 0.011 | 7.07 |
| SaaCH | 0.024 | 5.452 |
| VSA_EState 6 | 0.015 | 6.304 |

# CL

**Structural:** ZMIC2, ZMIC3, ZMIC4, Morg2, Morg3
**Electronic:** MATS1m, MATS4Z, MATS4m, AATS1dv, ATS6dv, ATS7dv, ATS7dv, ATS8dv, ATSC0are,
ATSC0pe, ATSC0se, ATS0dv, EState_VSA2, GATS3are, GATS3pe, GATS3se, GGI10

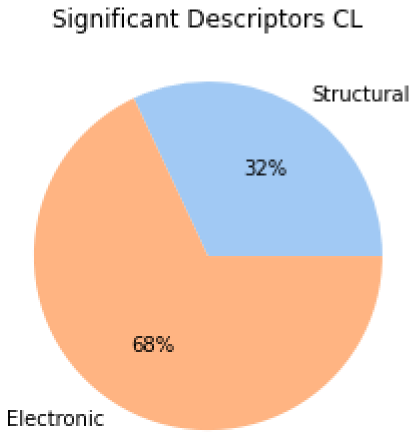


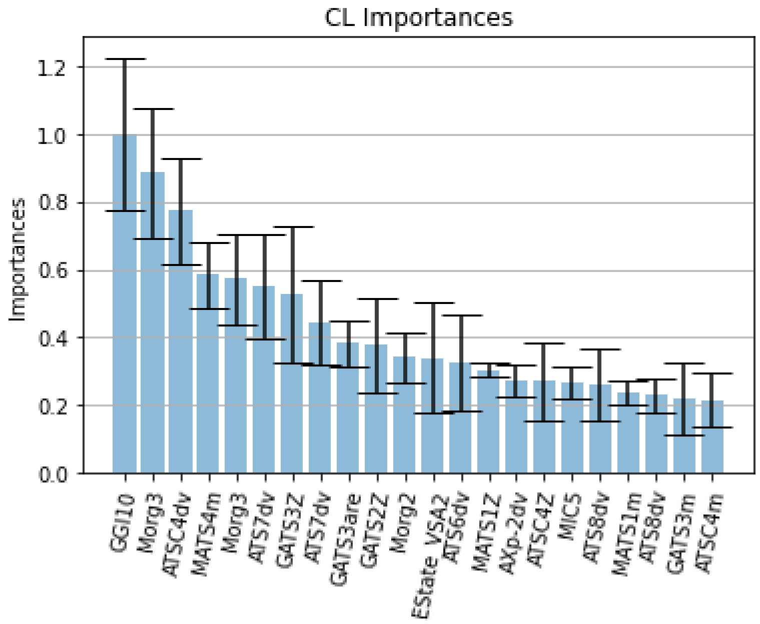


| Variable | P-score | F-score |
|---|---|---|
| ATSC4dv | 0.008 | 7.867 |
| MATS4m | 0.016 | 6.362 |
| MATS4Z | 0.015 | 6.523 |
| Morg2 | 0.012 | 6.988 |
| Morg3 | 0.009 | 7.59 |
| ATS8dv | 0.023 | 5.664 |

## CL2

**Structural:** nSpiro, C4SP3
**Electronic:** ATSC7c, ATSC8Z, ATSC8are, ATSC8dv, ATSC8i, ATSC8m, ATSC8pe, ATSC8se

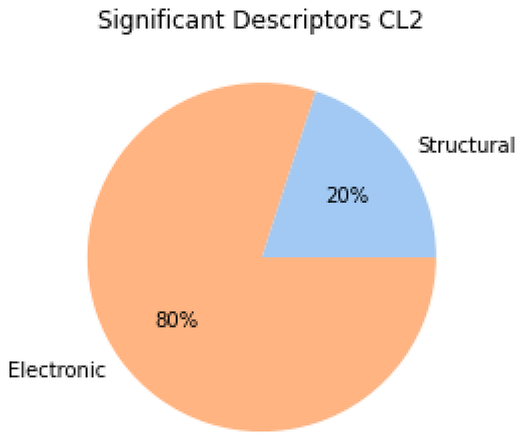


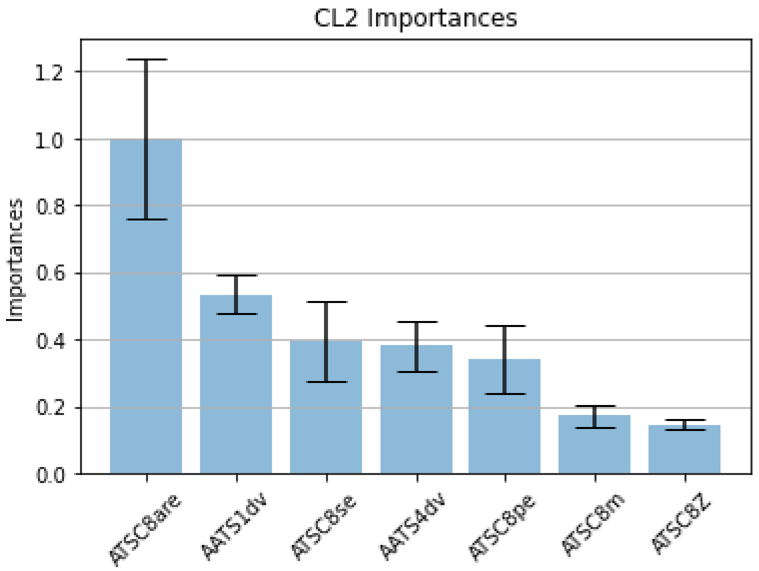


| Variable | P-score | F-score |
|---|---|---|
| C4SP3 | 0.004 | 9.244 |
| nSpiro | 0.004 | 9.349 |
| ATSC8i | 0.003 | 9.777 |
| ATSC7c | 0.012 | 6.861 |
| ATSC8Z | 0.008 | 7.618 |
| ATSC8pe | 0.001 | 12.23 |
| ATSC8m | 0.009 | 7.472 |
| ATSC8are | 0.001 | 11.386 |
| ATSC8se | 0.003 | 10.079 |
| ATSC8dv | 0.003 | 10.155 |

#### PST

**Structural:** NtsC, SsssB
**Electronic:** ATS5i, ATSC6are, ATSC6se, ATSC7c, ATSC7p, ATSC7v, ATS7are, ATS7i, ATSC8are, ATSC8se,
EState_VSA6

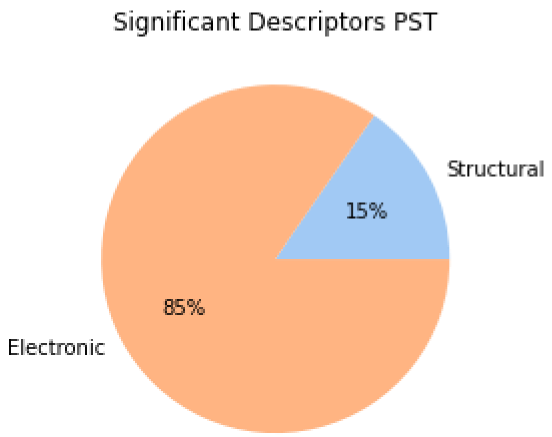


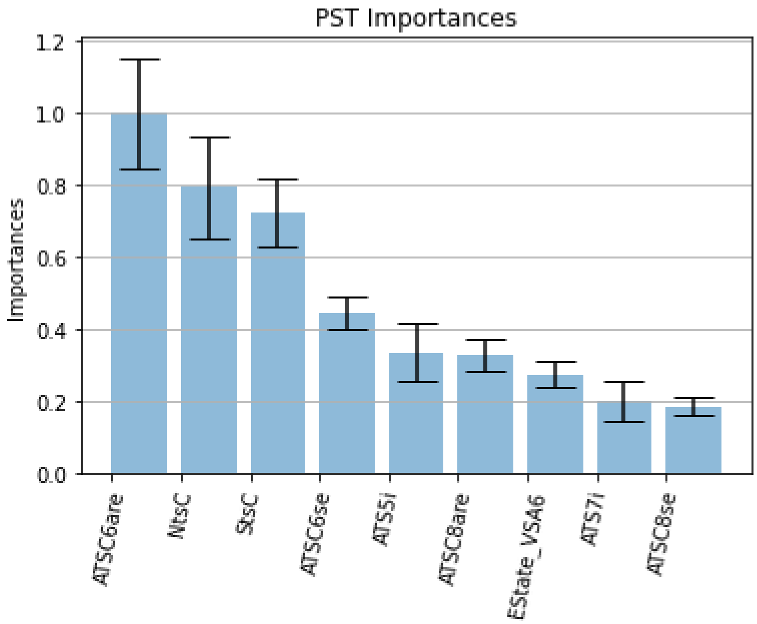


| Variable | P-score | F-score |
|---|---|---|
| ATSC7i | 0.006 | 8.426 |
| ATSC7p | 0.02 | 5.786 |
| ATSC7v | 0.019 | 5.91 |
| ATSC7c | 0.036 | 4.65 |
| ATSC7Z | 0.037 | 4.623 |
| ATSC7m | 0.042 | 4.371 |
| EState_VSA6 | 0.009 | 7.391 |

### PSB

**Structural:** nBase, SIC0, BIC0
**Electronic:** ATSC1dv, MATS1dv, GATS1d, GATS3d, VSA_EState9

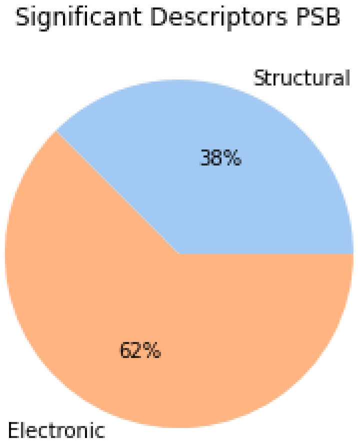


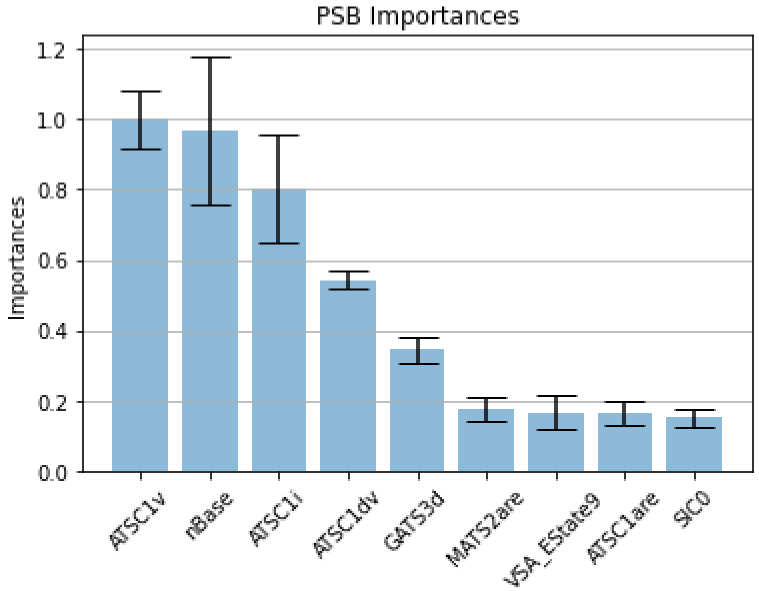


| Variable | P-score | F-score |
|---|---|---|
| SIC0 | 0.019 | 5.986 |
| BIC0 | 0.022 | 5.679 |
| GATS1d | 0.017 | 6.198 |
| nBase | 0.028 | 5.186 |
| VSA_EState9 | 0.029 | 5.138 |
| GATS3d | 0.027 | 5.259 |
| MATS1dv | 0.008 | 7.846 |
| ATSC1dv | 0.005 | 8.926 |
