## Supplementary Materials 3 for "An Artificial Intelligence Framework for Optimal Drug Design"

[LONGITUDINAL]
input = {KpISF, KpCSF, KpT, PSB, PST, CL, CL2, Ka, F}
F = {use = regressor}

PK:
depot(adm=1, target=Ap) ;#target=Ap for IV (adm 1)
depot(adm=2, target=Aa) ;#target=Aa for oral (adm 2)

; ### MODEL CODE ###

EQUATION:
	odeType = stiff

; ### PARAMETERS ###
;# Define Physiological Parameters
	;#Volumes (L)
	VBv = 0.0000502 ;#volume of brain plasma
	VCSF = 0.00012 ;#Volume of brain cerebrospinal fluid, 90-150uL
	VBev = 0.00228 - VBv - VCSF ;#Volume of brain extravasculature
	VTv = 0.00789468 ;#Volume of tissue vasculature
	Vp = 0.00906 - VBv - VTv ;#Volume of plasma in systemic circulation, excluding brain and tissue plasma
	VTev = 0.28-Vp-VBv-VCSF-VBev-VTv ;Volume of tissue extravasculature

	;# Flows (L/h)
	QCSF = 0.000132 ;#CSF flow to brain tissue and back from brain tissue
	QB = 0.0653 ;#Flow to brain vasculature
	QT = 2.8797 ;#Flow to tissue vasculature
	CLG = 0.00012 ;#Glymphatic clearance set to zero

	;#Permeability Surface Coefficients
	;#uncomment these lines for only one PSB parameter
	PSBBB = PSB * (0.0155/0.018) ;# (BBB_SA / Total_SA)
	PSCSF = PSB * (0.0025/0.018) ;#(BCSFB_SA / Total_SA)

	;#Combine KpISF and KpCSF into one KpB parameter
	;#uncomment these lines for only one KpB parameter
	;KpISF = KpB
	;KpCSF = KpB


;### INITIAL CONDITIONS ###
;#	

; ### EXPLICIT EQUATIONS ###

; ### DIFFERENTIAL EQUATIONS ###
ddt_Aa = -(Ka*F*Aa) - (Ka*(1-F)*Aa)

ddt_Ap = (QB*(CBv-Ap/Vp) + QT*(CTv-Ap/Vp) + CLG*CCSF - CL*Ap/Vp) + (Ka*F*Aa)

ddt_CBv = (QB*(Ap/Vp - CBv) + PSCSF*(CCSF/KpCSF - CBv) + PSBBB*(CBev/KpISF - CBv)) / VBv

ddt_CCSF = (PSCSF*(CBv-CCSF/KpCSF) + QCSF*((CBev/(KpISF/KpCSF)) - CCSF) - CLG*CCSF) / VCSF

ddt_CBev = (PSBBB*(CBv-CBev/KpISF) + QCSF*(CCSF - CBev/(KpISF/KpCSF))) / VBev

ddt_CTv = (QT*(Ap/Vp - CTv) + PST*(CTev/KpT - CTv)) / VTv

ddt_CTev = (PST * (CTv-CTev/KpT) - CTev*CL2) / VTev


BrainConc = (CBv*VBv + CBev*VBev + CCSF*VCSF) / (VBv + VCSF + VBev) ;# Concentration of Drug in Brain Homogenate, non-perfused

Cp = Ap/Vp

OUTPUT:
    output = {Cp, BrainConc, CBev, CCSF}
